# Pre-clinical efficacy of a C4BP hexameric IgG Fc fusion protein against *Neisseria gonorrhoeae*

**DOI:** 10.64898/2026.05.20.726552

**Authors:** Jutamas Shaughnessy, Jintang Du, Mary Wakim Broden, Sunita Gulati, Bo Zheng, Nancy Nowak, Gretchen Telford, Sandy P. Fontes, Y Tran, Keith L. Wycoff, Kevin J. Whaley, Alison K. Criss, Sanjay Ram

**Author notes:** **Correspondence:** Sanjay Ram, Division of Infectious Diseases and Immunology. Department of Medicine, University of Massachusetts Chan Medical School, Worcester, Massachusetts 01605, United States.

## Abstract

Gonorrhea is the second most common bacterial sexually transmitted infection and affects about 80 million people worldwide annually. The causative agent, *Neisseria gonorrhoeae*, has become resistant to almost every antibiotic used for its treatment. There is no licensed vaccine against gonorrhea. Therefore, there is an urgent need to develop novel prevention and treatment strategies to curb the spread of gonorrhea. The gonococcus has evolved several mechanisms to evade complement, a key arm of immune defenses against this pathogen, including binding of the human complement inhibitors Factor H (FH) and C4b-binding protein (C4BP). We previously showed that chimeric molecules fusing the gonococcal binding domains of FH and C4BP to IgG Fc and IgM Fc, respectively, mediate complement-dependent killing of gonococci in vitro and attenuate gonococcal colonization of mouse vaginas when administered topically. Here, we fused C4BP domains 1 and 2, which contain the gonococcal binding region, to IgG Fc bearing the IgM tail-piece to facilitate Fc hexamerization. This molecule, called C4BP-Hexa IgG Fc, showed ∼650-fold greater complement-dependent bactericidal activity on a molar basis than monomeric C4BP-IgG1 Fc. C4BP-Hexa IgG Fc enhanced association with and uptake by human neutrophils in a complement-independent manner. Despite ‘off-target’ complement activation in solution, C4BP-Hexa IgG Fc reduced both the duration and the bacterial burden of gonococcal vaginal colonization in human FH and C4BP transgenic mice when administered intravaginally daily. In conclusion, we show proof-of-concept of the efficacy of a hexameric C4BP IgG Fc fusion molecule against *N. gonorrhoeae*, which could aid in the fight against this multidrug-resistant pathogen.

## INTRODUCTION

*Neisseria gonorrhoeae* is the causative agent of gonorrhea, which affects about 80 million people globally annually (1). Gonorrhea primarily infects mucosal surfaces and causes urethritis in men and cervicitis in women. The pharynx, rectum and conjunctivae can be infected in both sexes. Serious complications in women include endometritis and salpingo-oophoritis, which can lead to sequelae such as tubal infertility, ectopic pregnancy and chronic pelvic pain. In some instances, the gonococcus can spread to distant tissues through the bloodstream to cause disseminated gonococcal infection (DGI). Manifestations of DGI include tenosynovitis, skin lesions and septic arthritis (2, 3). In rare instances, it can infect heart valves to cause endocarditis or invade the central nervous system to cause meningitis.

*N. gonorrhoeae* has become resistant to almost every antibiotic that has been used for its treatment (4, 5). Currently, ceftriaxone is the only recommended first-line treatment for gonorrhea (6). Alarmingly, resistance to ceftriaxone has been reported from several countries (7-11), with rates approaching 30% in certain countries in South-East Asia (12, 13). Two new oral topoisomerase IV inhibitor antibiotics, zoliflodacin (a spiropyrimidinetrione) (14) and gepotidacin (a triazaacenaphthylene) (15), which target type II topoisomerases (the GyrA subunit of DNA gyrase and the ParC subunit of topoisomerase IV), have been approved by the FDA for the treatment of uncomplicated gonococcal infections (16). However, treatment failures and rising MICs of both antibiotics have been reported (17-19). Currently, there are no licensed vaccines against gonorrhea. Retrospective analyses have shown that a group B meningococcal vaccine containing detergent-extracted outer membrane vesicles is associated with a 30-40% reduction in gonorrhea rates (20-24). However, these vaccines may have limited efficacy in persons with chlamydia (20, 23) and in high-risk men who have sex with men (MSM) (25, 26). Thus, novel therapies to combat gonorrhea are urgently needed.

Complement is an important arm of innate immune defenses against gonorrhea (27-29). Gonococci have evolved several mechanisms to evade complement, including binding of two complement inhibitors: Factor H (FH), an inhibitor of the alternative pathway, and C4b-binding protein (C4BP), an inhibitor of the classical pathway (30-32). We previously showed that fusing the gonococcal binding domains of FH to the Fc domain of human IgG to create an FH-Fc chimeric molecule was efficacious in the mouse vaginal colonization model of gonorrhea when given topically intravaginally (33-36). Similarly, fusing the N-terminal two domains of the α-chain of C4BP that harbors the gonococcal binding site, but lacks any complement regulating activity, to the Fc domain of IgM was also effective against gonococci in mice when administered intravaginally (37).

IgM Fc was chosen for the C4BP-Fc chimeric protein to create a multimeric (10 or 12 C4BP ‘arms’) molecule to effectively out-compete the binding of endogenous C4BP, which contains either six or seven α-chains (38). However, manufacture of IgM at scale is technically challenging. Introduction of the IgM Fc tail-piece at the C-terminus of IgG Fc results in the formation of IgG Fc hexamers (39), which permits high-avidity binding while facilitating purification through traditional Protein A-based affinity chromatography. Like IgM, a single hexameric IgG molecule can engage the C1 complex and activate the classical pathway, resulting in greater complement-dependent cytotoxicity compared to conventional monomeric IgG. Accordingly, a hexameric IgG derivative of rituximab, a CD20-targeting monoclonal antibody (mAb), showed significantly enhanced complement-dependent cytotoxicity (CDC) (39). Here, we evaluated the efficacy of a chimeric protein comprising C4BP domains 1 and 2 fused to hexameric human IgG1 Fc (C4BP-Hexa IgG Fc) against *N. gonorrhoeae*.

## RESULTS

### Binding of C4BP-Hexa IgG Fc to N. gonorrhoeae

Binding of C4BP-Hexa IgG Fc to *N. gonorrhoeae* was measured by flow cytometry (**Fig. 1**). Dose-dependent binding of C4BP-Hexa IgG Fc to FA1090 and MS11 was observed, both of which bind human C4BP (30). By contrast, no binding was observed to strains H041 and F62, which do not bind human C4BP (30, 33).

**Fig. 1.**
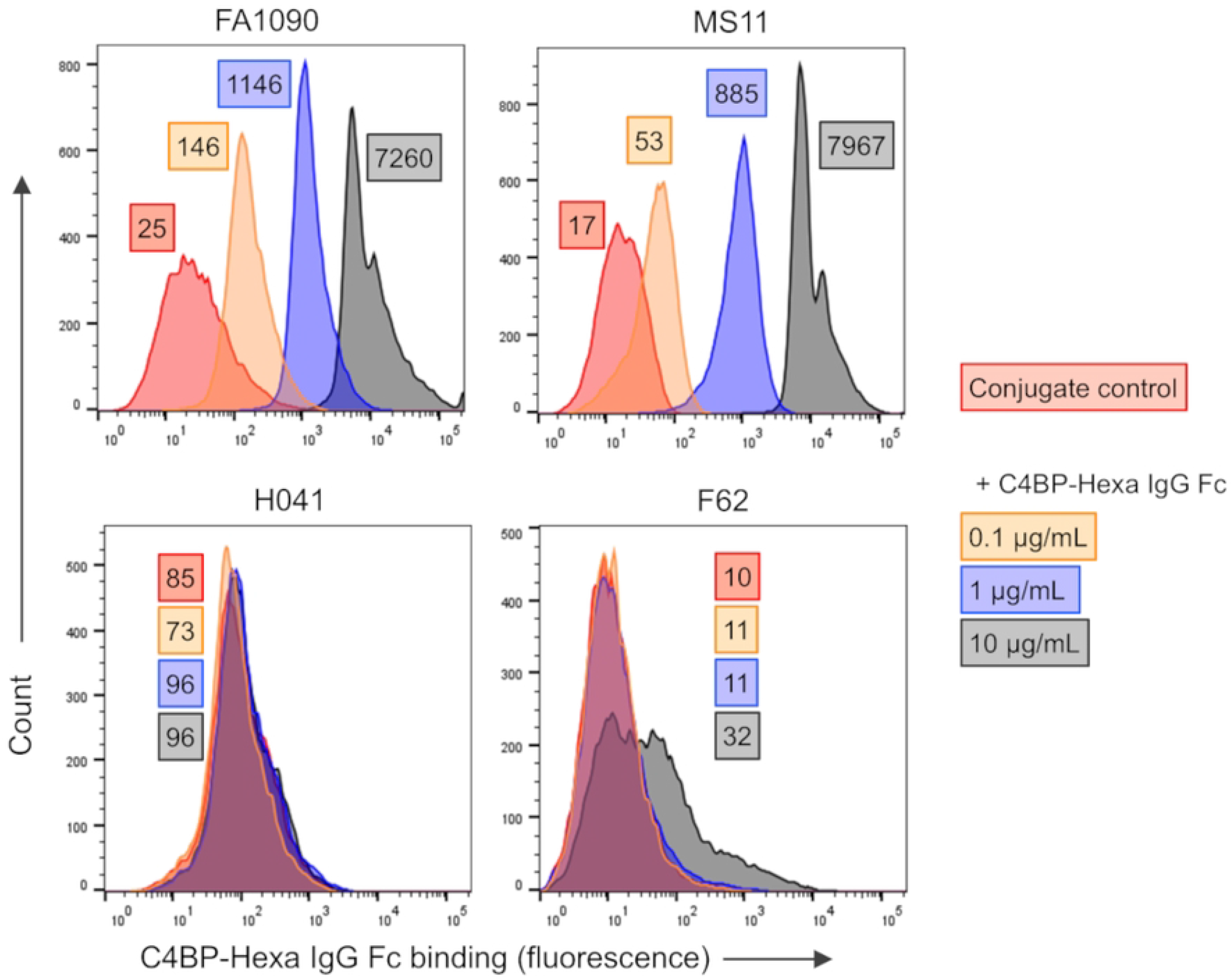
Binding of C4BP-Hexa IgG Fc to *N. gonorrhoeae*. Gonococcal strains FA1090, MS11 and H041 (WHO X) were incubated with 0.1, 1 or 10 µg/mL of C4BP-Hexa IgG Fc, or with buffer alone (conjugate control). Bound C4BP-Hexa IgG Fc was detected with anti-human IgG FITC. X-axis, fluorescence; Y-axis, counts. One representative experiment of two independent experiments is shown.

### C4BP-Hexa IgG Fc mediates complement-dependent bactericidal activity of N. gonorrhoeae

The ability of C4BP-Hexa IgG Fc to mediate complement-dependent killing of *N. gonorrhoeae* was measured in serum bactericidal assays using IgG- and IgM-depleted normal human serum (NHS) as the complement source. As shown in **Fig. 2**, C4BP-Hexa IgG Fc was highly bactericidal against strains MS11 and FA1090, with absolute IC_50_ values (the concentration of fusion protein calculated to yield 50% killing (50% survival)) of 0.009 µg/mL and 0.091 µg/mL, respectively. As expected, no killing of C4BP non-binding strains H041 and F62 was observed. In comparison, the IC_50_ of C4BP-IgG1 Fc (monomeric Fc) against FA1090 was 9.03 µg/mL, which was about 100-fold higher than the IC_50_ of C4BP-Hexa IgG Fc versus FA1090. On a molar basis, this translates to an approximately 650-fold greater activity of the hexameric molecule compared to the monomeric Fc fusion protein.

**Fig. 2.**
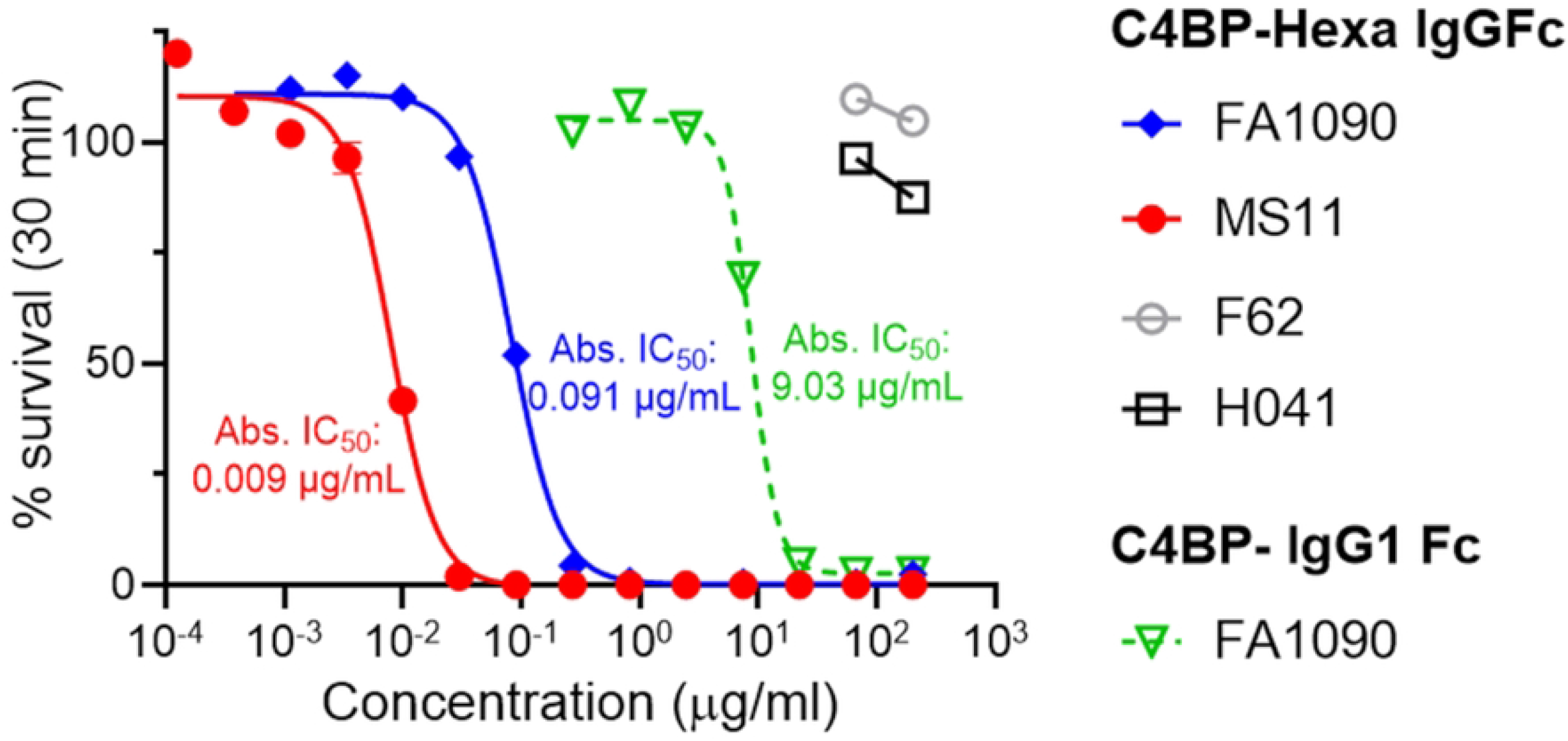
Bactericidal activity of C4BP-Hexa IgG Fc. *N. gonorrhoeae* strains FA1090 and MS11 (C4BP binders) and H041 and F62 (C4BP non-binders) were incubated with the indicated concentrations of C4BP-Hexa IgG Fc, followed by the addition of human complement (IgG and IgM depleted normal human serum) to a final concentration of 10% (v/v). The efficacy of C4BP-IgG1 Fc (monomeric Fc) against FA1090 was also measured. Survival of bacteria following 30 min of incubation relative to 0 min, expressed as a percentage, is indicated on the Y-axis. A sigmoidal 4-parameter logistic curve was plotted to determine the absolute IC_50_ (Abs. IC_50_), which is the calculated concentration of C4BP-Hexa IgG Fc or C4BP-IgG1 Fc predicted to yield 50% killing (50% survival). IC50s are indicated to the right of the curves; the color of the font corresponds to the color of the curves.

### C4BP-Hexa IgG Fc enhances association and uptake of gonococci by human neutrophils

We next asked whether C4BP-Hexa IgG Fc could enhance gonococcal association with and uptake by neutrophils (polymorphonuclear leukocytes, PMNs). For these assays, we used a FA1090 mutant where all Opacity protein genes *(opa)* were deleted (FA1090 Opa-) (40) to prevent confounding effects of nonopsonic, Opa-mediated uptake by PMN carcinoembryonic antigen-related adhesion molecule 3 (CEACAM3) (41, 42). We compared the abilities of C4BP-Hexa IgG Fc, C4BP-IgG Fc (C4BP domains 1 and 2 fused to monomeric human IgG1 Fc) and polyclonal rabbit anti-gonococcal IgG to mediate attachment (association) to and uptake of gonococci by PMNs. These experiments were performed in the absence of complement. As shown in **Fig. 3**, C4BP-Hexa IgG Fc, C4BP-IgG Fc and rabbit anti-gonococcal IgG all facilitated the association and uptake of gonococci by PMNs to a similar extent in a complement-independent manner.

**Fig. 3.**
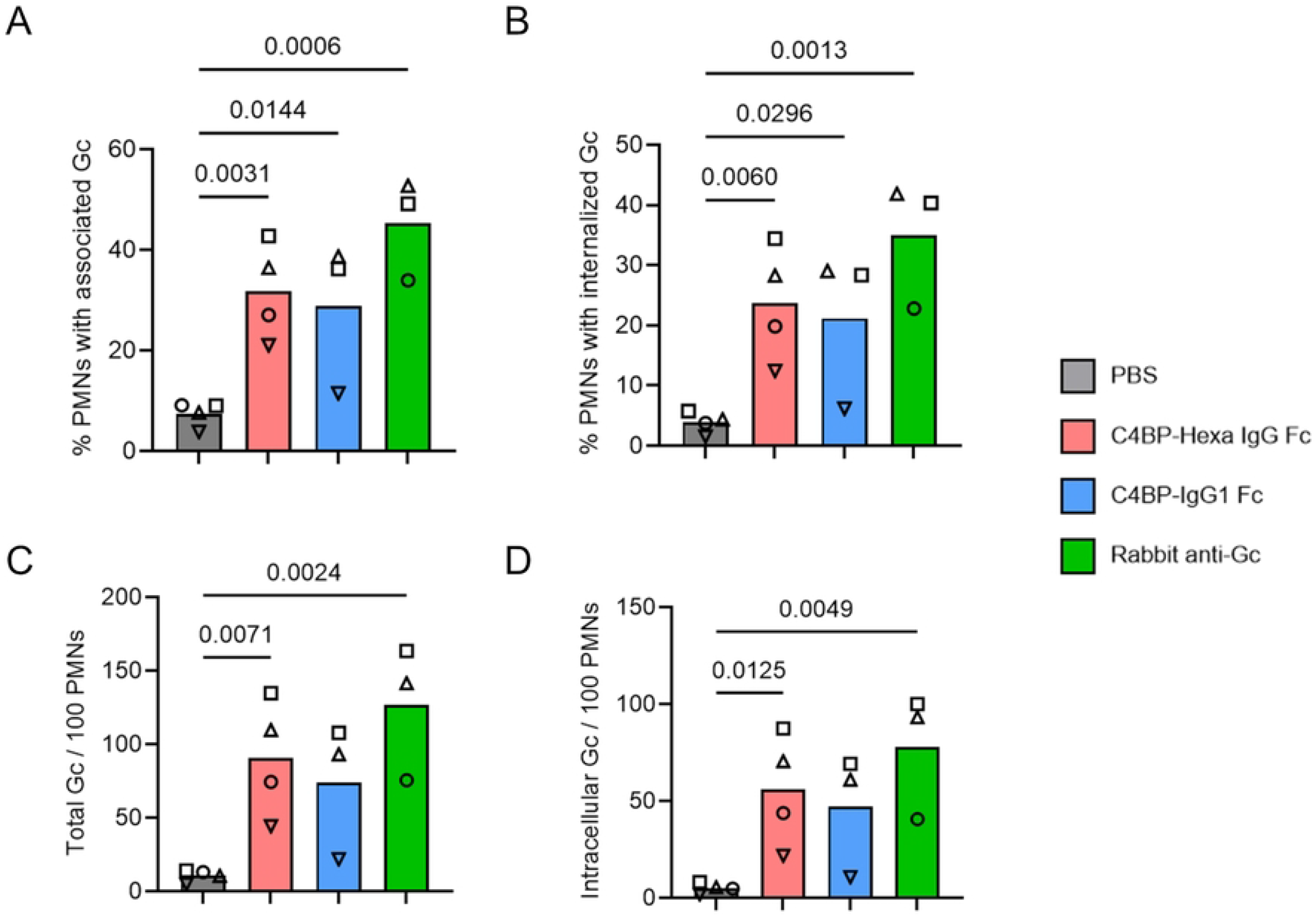
C4BP-Hexa IgG-Fc enhances opsonophagocytosis of *N. gonorrhoeae*. An *N. gonorrhoeae* FA1090 mutant where all genes encoding for its opacity proteins (Opa) were deleted (FA1090 Opa-) was incubated with C4BP-Hexa IgG Fc (30 µg/mL), C4BP-IgG1 Fc (C4BP domains 1 and 2 fused to a monomeric human IgG1 Fc, 100 µg/mL), polyclonal rabbit anti-N. gonorrhoeae IgG (Rabbit anti-Gc at 500 µg/mL that was used as a positive control) or PBS (negative control) and PMNs purified from human blood. The multiplicity of infection (MOI; ratio of gonococci to PMNs) was 1. The mixture was incubated for 1 h at 37 °C. **A**. Percentage of PMNs associated with gonococci (Gc). **B**. Percentage of PMNs with internalized Gc. **C**. Number of Gc associated with PMNs (Total number of Gc per 100 PMNs). **D**. Number of intracellular Gc per 100 PMNs. The mean of 3 – 4 separate experiments is shown. Each experimental replicate is designated by a different shape. Comparisons across groups were made by the mixed-effect analysis and pairwise comparisons across groups by Tukey’s multiple comparisons test.

### C4BP-Hexa IgG Fc activates the classical complement pathway in solution

Unlike IgM, hexameric forms of IgG, formed either through covalent linking via the C-terminal IgM tailpiece (39) or through Fc mutations that drive non-covalent Fc-Fc interactions and hexamer formation in solution (43), activate complement in solution (off-target complement activation). We determined classical pathway activation in solution by adding increasing amounts of C4BP-Hexa IgG Fc to normal human serum (NHS) and measuring C4d generation. As shown in **Fig. 4**, C4BP-Hexa IgG Fc at a concentration of 10 µg/mL in serum generated 2.7-fold more C4d compared with NHS containing 10 µg/mL of heat-aggregated IVIg. Reducing the concentration of C4BP-Hexa IgG Fc to 1 µg/mL reduced C4d generation to baseline levels seen with NHS alone.

**Fig. 4.**
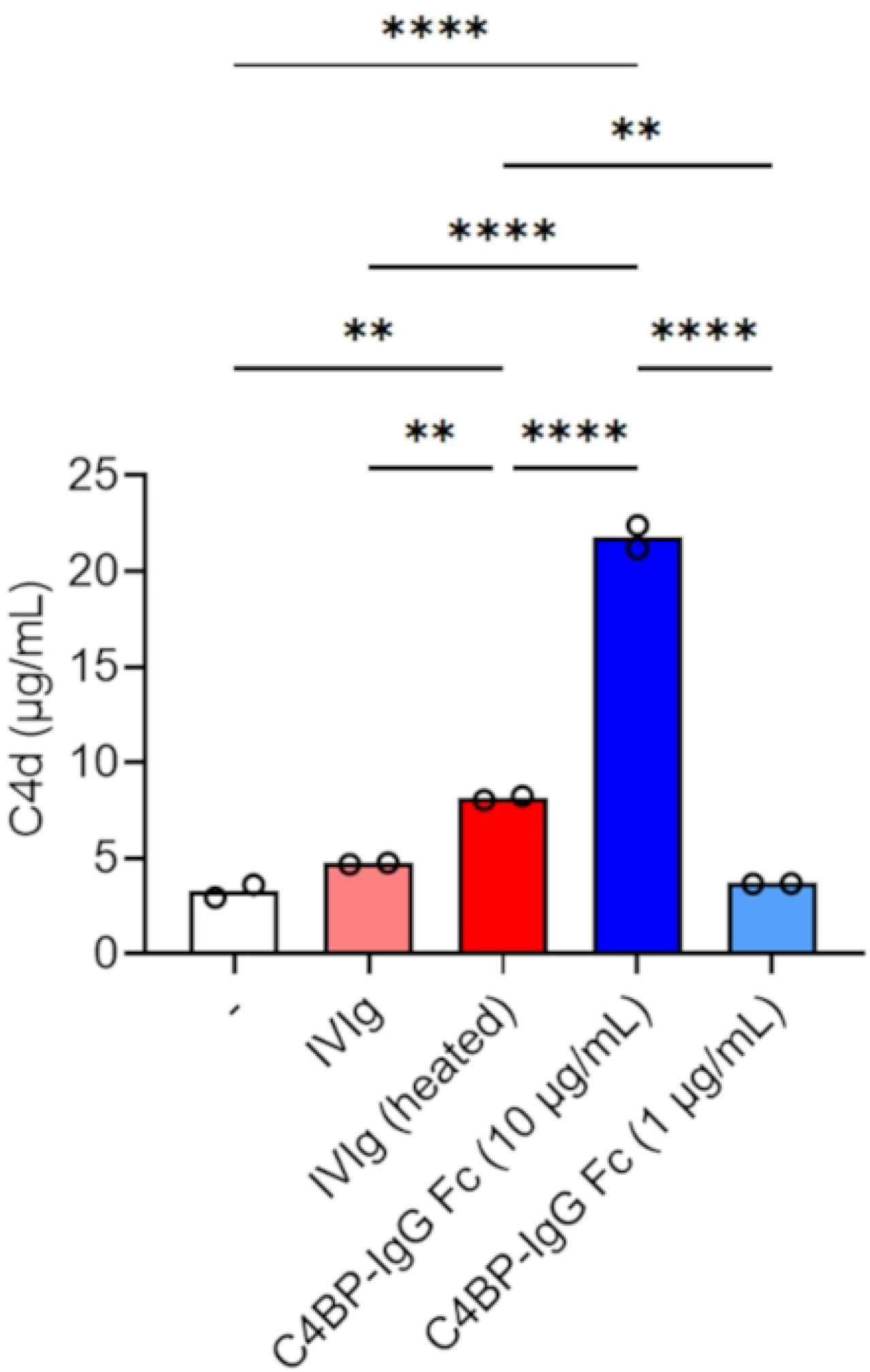
C4BP-Hexa IgG Fc activates the classical pathway of complement in solution. Normal human serum (NHS) was incubated with either PBS (baseline C4d generation, indicated by “-”), soluble Intravenous immunoglobulin (IVIg (Gammagard), 10 µg/mL), heat-aggregated IVIg (10 µg/mL) or C4BP-Hexa IgG Fc (10 or 1 µg/mL) at 37 °C for 60 min. C4d generated was quantified by ELISA using the MICROVUE C4d ELISA kit (QuidelOrtho). Data from two independent experiments are shown. Pairwise comparisons across groups were made by repeated measures one-way ANOVA with Tukey’s multiple comparisons test. **, P<0.01; ***, P<0.001; ****, P<0.0001.

### Efficacy of topically administered C4BP-Hexa IgG Fc in the mouse vaginal colonization model of gonorrhea

We next assessed the efficacy of C4BP-Hexa IgG Fc in the mouse vaginal colonization model of gonorrhea using transgenic mice that expressed two human complement inhibitors, Factor H (FH) and C4BP (FH/C4BP Tg mice) (33, 36, 44). Gonococci bind only human, but not mouse, FH and C4BP (45, 46). Therefore, we used FH/C4BP Tg mice to provide the colonizing gonococci the complement evasion mechanisms available to them in humans. In contrast to FH/C4BP Tg mice, complement activation mediated by C4BP-Hexa IgG Fc would be unimpeded in wild-type mice, which may spuriously enhance its therapeutic efficacy. As shown in **Fig. 5A-C**, C4BP-Hexa IgG Fc significantly reduced the duration and burden of colonization of strain FA1090. However, there was no reduction of the duration or burden of colonization with strain F62, which does not bind C4BP (**Fig. 5D-F**).

**Fig. 5.**
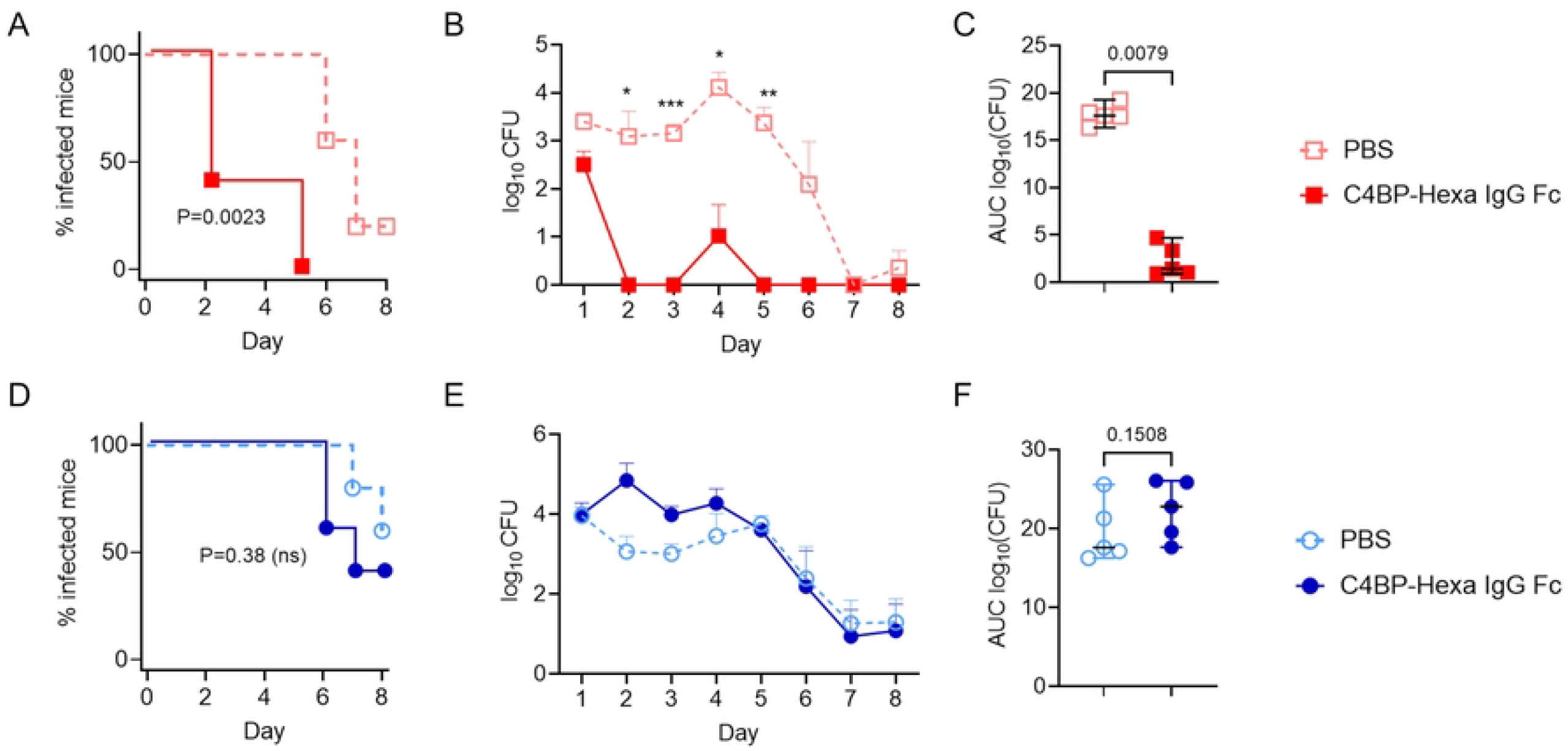
Efficacy of C4BP-Hexa IgG Fc in the mouse vaginal colonization model of gonorrhea. Eight-week-old human Factor H and C4b-binding protein (C4BP) transgenic (FH/C4BP Tg) mice in the diestrus phase of the estrus cycle were treated with antibiotics and 17β-estradiol. Mice were colonized intravaginally on day 0 with 3.6 × 10^7^ CFU FA1090 (top panel, **A-C**) or 2.8 × 10^7^ CFU F62 (bottom panel, **D-F**). Starting on day 1, mice (n=5 per group) were given 5 µg of C4BP-Hexa IgG Fc in 10 µL of PBS, or 10 µL of PBS alone (vehicle control) daily through day 8. Vaginas were swabbed daily to enumerate CFUs. **A** and **D**: Kaplan Meier curves showing time to clearance. Comparisons between groups were made by Mantel-Cox log-rank test. **B** and **E:** log_10_ CFU versus time. Comparisons between groups were made by two-way ANOVA and Šídák’s multiple comparisons test. *, P<0.05; **, P<0.01; ***, P<0.001. **C** and **F:** Area Under Curve (AUC) analysis. The mean and 95% confidence intervals are shown. The two groups were compared by the Mann-Whitney non-parametric test.

## DISCUSSION

In this study we demonstrate the efficacy of a hexameric IgG1 Fc construct fused to the two N-terminal domains of C4BP against *N. gonorrhoeae* in vitro and in the mouse vaginal colonization model. This study mirrors prior work where a construct with the same two C4BP domains fused to IgM Fc was bactericidal and attenuated colonization of mouse vaginas (37). Both multimeric fusion proteins are potent complement activators and effectively overcome the complement inhibition mediated by endogenous C4BP. IgM-based therapeutics have been tested in human trials, but currently none have been FDA approved (47). While IgM therapeutics face manufacturing challenges, molecules bearing IgG Fc can be purified using well-established methods, including Protein A chromatography and size-exclusion chromatography.

Prior work has shown that the mechanism of action of a FH-Fc fusion protein that comprised the two C-terminal domains of FH relied on an intact terminal complement pathway, suggesting that gonococcal clearance was mediated through membrane attack complex (MAC, or C5b-9) (36). The female genital tract contains hemolytically active complement (48), which can support the killing of gonococci by Fc fusion proteins and an anti-LOS monoclonal antibody (36, 49). While the likely mechanism of action of C4BP-Hexa IgG Fc also involves MAC, C4BP-Hexa IgG Fc also enhances the uptake of gonococci by PMNs independent of complement, which could also contribute to gonococcal clearance.

Despite its ability to activate complement in solution, C4BP-Hexa IgG Fc was efficacious *in vivo*, suggesting sufficient active complement was available to mediate gonococcal killing. When designing a sustained-release formulation of C4BP-Hexa IgG Fc, it would be important to ensure sufficient concentrations for therapeutic efficacy, yet not exceed amounts that may result in complement consumption in vivo. The kidney, endothelium and red blood cells are highly susceptible to complement-mediated damage. Given its ability to activate complement in solution, caution should be exercised if C4BP-Hexa IgG Fc is given systemically. Sopp et al showed that introducing the C575S mutation in the IgM tail-piece prevents covalent IgG hexamer formation through disulfide bonds and ‘off-target’ complement activation in solution, but results in enhanced IgG hexamerization following binding to surfaces (‘on-target hexamerization’) (39). While complement-dependent cytotoxicity mediated by a C575S variant of the anti-CD20 mAb rituximab was greater than that of rituximab with wild-type IgG1 Fc, it was less than that of the Hexa IgG rituximab (i.e., bearing the native IgM tail-piece that forms covalent Fc hexamers) (39). C4BP-Hexa IgG Fc has the advantage of multivalent high-avidity binding, which can effectively out-compete endogenous C4BP.

In a previous study, binding of a chimeric protein that comprises C4BP domains 1,2 and 4 through 8 (i.e., all α chain domains with the exception of domain 3) fused to IgG1 Fc, but not a C4BP domain 1-2 IgM Fc protein, was competed out by native C4BP. We concluded that multivalent binding of C4BP was required to compete with binding of endogenous human C4BP, which prompted evaluation of the therapeutic potential of the IgM Fc fusion protein. In accordance with the prior study, we noted that the hexameric Fc molecule had an ∼650-fold increase in activity on a molar basis compared to the monomeric molecule. Whether a bivalent C4BP-IgG Fc molecule that has the capacity to hexamerize following binding (i.e., ‘on-target’ hexamerization), as may be achieved with mutations such as E430G (43), the triple Q311R/M428E/N434W (collectively called the REW mutations) (50), or with an IgM tail-piece bearing the C575S mutation (39), and mediate killing remains to be determined.

In a large strain survey, Bettoni et al found that 96/107 (89.7%) of PorB1a isolates, but only 16/83 (19.3%) of PorB1b isolates bound C4BP (37). Preferential binding of C4BP to PorB1a-expressing isolates has been demonstrated in other studies (30, 33). Thus, efficacy of C4BP-Hexa IgG Fc will be largely limited to PorB1a-expressing strains, which cause 10% – 30% of all gonococcal infection in most series (51, 52). However, PorB1a-expressing strains are about 20 times more likely to disseminate than their PorB1b counterparts (52). Disseminated gonococcal infection (DGI) can cause considerable morbidity, and complications such as endocarditis and meningitis can be fatal. Therefore, coverage of PorB1a strains is likely to have a profound impact in reducing the incidence of DGI. We previously showed that a chimeric protein comprising IgG3 Fc fused to FH domains 19 and 20 supported complement-dependent bactericidal activity of all 46 PorB1b strains tested, but only 2 of 15 PorB1a strains (33). Thus, combining FH-Fc with a C4BP-Fc chimeric would provide broad activity against almost all gonococcal isolates.

Similar to FH-Fc, C4BP-Hexa IgG Fc targets another key complement evasion mechanism. Resistance to C4BP-Hexa IgG Fc would require PorB mutations that abrogate C4BP binding, which would render the organism susceptible to complement and thereby reduce fitness in vivo. In conclusion, we provide proof-of-concept for the efficacy of C4BP-Hexa IgG Fc against C4BP-binding *N. gonorrhoeae*, which targets a critical virulence mechanism and could be a valuable tool to combat the spread of gonorrhea.

## MATERIALS AND METHODS

### Expression and purification of C4BP-hexameric IgG Fc fusion protein

The manufacture of C4BP-hexameric IgG was conducted at KBio/ZabBio labs (San Diego, CA). To generate C4BP-Hexa IgG Fc, DNA sequences coding for CCP1-2 domains of C4BP and human IgG1-Fc were cloned in frame with N-terminal signal peptide sequence and C-terminal IgM tail piece into the viral based expression vector (pJL TRBO) (37, 53, 54). DNA fragments were ordered from Thermo Fisher with GeneArt codon optimization for *Nicotiana benthamiana. N. benthamiana* variety deficient in xylosyl- and fucosyltransferase was vacuum infiltrated with agrobacterium strain carrying gene of interest (C4BP hexamer) into whole plants (55). Seven days post-infiltration, leaf material was harvested, homogenized, and filtered using miracloth (Millipore). C4BP-Hexa IgG Fc was purified via protein A chromatography, then Capto core 400 chromatography was used to separate hexameric and monomeric forms. Size and purity were verified by SDS-PAGE. C4BP-IgG1 Fc was newly generated for this study. The nucleotide sequences encoding (from 5’ to 3’) human C4BP CCP domains 1 and 2 fused to the IgG1 hinge constant portion of human IgG1 was synthesized and cloned into mammalian expression vector pcDNA 3.4 by GenScript (Piscataway, NJ). Expi-CHO cells (ThermoFisher) were transfected with the plasmid and the fusion protein, called C4BP-IgG1 Fc, was purified from tissue culture supernatants using affinity chromatography over Protein A/G agarose. Purity and size of the obtained proteins were verified by SDS-PAGE. Protein concentration was quantified by absorption at 280 nm.

### N. gonorrhoeae strains

Strains FA1090 (PorB1b) (56), MS11 (PorB1b) (57), H041 (WHO X; PorB1b) (8) and 15253 (PorB1a) (58) have been described previously. Studies with PMNs used FA1090 1-81-S2 Opa-negative (Opa-) background, in which all 11 *opa* genes are chromosomally deleted, has been described previously (40).

### Binding of C4BP-hexa IgG Fc by flow cytometry

Binding of C4BP-Hexa IgG Fc to *N. gonorrhoeae* was performed by flow cytometry as described previously (33). Briefly, 10^7^ CFU *N. gonorrhoeae* suspended in HBSS containing 1 mM CaCl_2_ and 1 mM MgCl_2_ (HBSS^++^) were incubated with the indicated concentrations of C4BP-Hexa IgG Fc for 30 min at 37 °C. Bacteria were washed twice in HBSS^++^ and bound C4BP-Hexa IgG Fc was detected with anti-human IgG FITC (Sigma). Bacteria were washed and fixed with PBS containing 1% paraformaldehyde, binding (fluorescence) was measured on a BD LSR II flow cytometer, Data were analyzed using FlowJo software.

### Serum bactericidal assay

Serum bactericidal assays were performed using a modification of previously described methods (59). Briefly, ∼1000 CFU *N. gonorrhoeae* were incubated with the indicated concentrations of C4BP-Hexa IgG Fc, followed by the addition of human complement (IgG and IgM depleted normal human serum (NHS); Pel-Freez) to a final concentration of 10 %. The final volume of the reaction mixture was 50 µL. Aliquots of bacteria were plated on chocolate agar at 0 and 30 min. Survival at 30 min relative to colony forming unit (CFU) at 0 min was expressed as a percentage.

### Primary human neutrophil isolation

Venous blood from healthy human donors was collected following obtaining written informed consent in accordance with a protocol approved by the University of Virginia Institutional Review Board for Health Science Research (protocol #13909). Subjects were recruited in January and February, 2026. Neutrophils were isolated as previously described (60), using dextran sedimentation, Ficoll gradient centrifugation, and red blood cell lysis.

### Imaging flow cytometry for association and internalization of gonococci by primary human neutrophils

Adherent, IL-8 treated primary human neutrophils were infected with Tag-it-Violet™ (TIV)-labeled Gc that were incubated with C4BP-hexa IgG Fc (30 µg/mL), C4BP-IgG1 Fc (100 µg/mL) or polyclonal rabbit anti-gonococcal (500 µg/mL) as a positive control (61). Incubation of bacteria with opsonins was done in Dulbecco’s PBS containing 0.9 mM CaCl_2_ and 0.5 mM MgCl_2_, and infection of PMNs was done in RPMI containing 10% heat-inactivated FBS. Bacterial internalization was assessed by imaging flow cytometry as previously described (62-64). Extracellular Gc was identified via staining with rabbit anti-Gc antibody (Meridian B65111R) coupled to DyLight (DL) 650 (Thermo) in-house. Intracellular bacteria were identified as TIV(+) DL650(-) particles associated with focused, single neutrophils, and results are expressed as the percentage of neutrophils containing at least one intracellular gonococcus. Neutrophils containing at least one associated bacterium and the number of total and intracellular gonococci per 100 neutrophils were also quantified.

### C4d ELISA

C4d generated by the addition of C4BP-Hexa IgG Fc to human complement was measured by the MICROVUE C4d ELISA kit (QuidelOrtho). Increasing amounts of C4BP-Hexa IgG Fc were added to human complement, and the mixture was incubated at 37 °C for 1 h. A positive control consisted of IgG (IVIg; Gammagard) aggregated by heating at 65 °C for 1 h (65). The assay was performed according to the manufacturer’s instructions.

### Mouse vaginal colonization model

Use of animals in this study was performed in strict accordance with the recommendations in the Guide for the Care and Use of Laboratory Animals by the National Institutes of Health. The protocol (PROTO202000074) was approved by the Institutional Animal Care and Use Committee at the University of Massachusetts Chan Medical School. Female mice 6–8 weeks of age in the diestrus phase of the estrous cycle were implanted with slow-release estrogen (*i*.*e*., 17β-estradiol) pellet (5 mg; Innovative Research of America, Cat. No. E-121) on day -2 prior to *N. gonorrhoeae* inoculation as described previously (66). Antibiotics (vancomycin, trimethoprim, and streptomycin) ineffective against *N. gonorrhoeae* were used to reduce competitive microflora (67). Mice were infected (day 0) with *N. gonorrhoeae* (strain and inoculum specified for each experiment), and vaginal swabs were performed daily on individual mice. *N. gonorrhoeae* CFUs in individual vaginal samples were determined by serial dilution and plating on chocolate agar supplemented with vancomycin, colistin, nystatin, trimethoprim and streptomycin (VCNTS; Thermo Fisher Scientific Inc., Cat. B12408).

### Statistical analyses

A sigmoidal 4-parameter logistic curve was plotted for killing of bacteria by various concentration of C4BP-Hexa IgG Fc. The IC_50_, defined as the concentration of mAb expected to yield 50% killing (i.e., 50% survival), was calculated by interpolation onto the sigmoidal 4PL curve. Comparisons across groups in the opsonophagocytosis and C4d ELISA assays were performed by one-way ANOVA and pairwise comparisons by Tukey’s post-test. Three parameters of efficacy were measured in the mouse infection experiments: time to clearance was plotted using Kaplan-Meier curves and the groups were compared by Mantel-Cox log-rank test; log_10_ CFU vs time, where comparisons across groups were made by two-way ANOVA and Šídák’s multiple comparisons test, and Area Under Curve (AUC) analysis, where the log_10_ AUC of each mouse was determined and the means of the two groups was compared by the Mann-Whitney test. All statistics were performed using GraphPad Prism 10.

## Notes

### Competing Interest Statement

The authors have declared no competing interest.

